# Performance Scaling for Structural MRI Surface Parcellations: A Machine Learning Analysis in the ABCD Study

**DOI:** 10.1101/2021.10.18.464804

**Authors:** Sage Hahn, Max M. Owens, DeKang Yuan, Anthony C Juliano, Alexandra Potter, Hugh Garavan, Nicholas Allgaier

## Abstract

The use of pre-defined parcellations on surface-based representations of the brain as a method for data reduction is common across neuroimaging studies. In particular, prediction-based studies typically employ parcellation-driven summaries of brain measures as input to predictive algorithms, but the choice of parcellation and its influence on performance is often ignored. Here we employed pre-processed structural magnetic resonance imaging data (sMRI) from the ABCD Study^®^ to examine the relationship between 220 parcellations and out-of-sample predictive performance across 45 phenotypic measures in a large sample of 9-10-year-old children (N=9,432). Choice of Machine Learning (ML) pipeline and use of alternative multiple parcellation-based strategies were also assessed. Relative parcellation performance was dependent on the spatial resolution of the parcellation, with larger numbers of parcels (up to ∼4000) outperforming coarser parcellations, according to a power-law scaling of between 1/4 and 1/3. Performance was further influenced by the type of parcellation, ML pipeline, and general strategy, with existing literature-based parcellations, a support vector based pipeline, and ensembling across multiple parcellations, respectively, as the highest performing. These findings highlight the choice of parcellation as an important influence on downstream predictive performance, showing in some cases that switching to a higher resolution parcellation can yield a relatively large boost to performance.

## Introduction

The application of Machine Learning (ML) methodologies to associate structural magnetic resonance imaging (sMRI) measures with phenotypic variation is an increasingly popular approach for studying brain function and structure (Davatzikos 2019). Further, working with surface-level representations of sMRI is a common technique employed by a number of popular neuroimaging software packages, such as FreeSurfer (Fischl 2012). For example, a common surface-based ML workflow is to use FreeSurfer-derived ROIs, either alone or with other features, to predict a phenotype of interest (Boeke 2020, Sato 2013, Hong 2020, Bhagwat 2019, Hahn 2020, He 2020). Despite the ubiquity of this and similar approaches, the degree to which the choice of parcellation might affect downstream performance is less well studied.

### Parcellations

Brain parcellations are used across various neuroimaging modalities. At their core, they represent a method for reducing the dimensionality of a large three-dimensional volume or surface representation of the brain to a lower dimensional representation (Eickhoff, 2017).

Parcellations typically have the added benefit of providing increased interpretability compared to other approaches to dimension reduction by their explicit reduction to spatially contiguous regions or networks of interest. Regions in this case define a group of voxels (3D equivalent to a pixel / unit) or vertices (surface / network base unit), with each region constituting a distinct spatial area (athough some parcellations allow overlaps between regions); networks define a collection of not necessarily spatially contiguous regions. Full (i.e., unparcellated) volumetric or surface-based brain data typically requires specifically designed analytic tools, as they contain tens to hundreds of thousands of variables per scan. In contrast, parcellations allow researchers to more easily employ traditional statistical or ML based methods in studying brain structure and function. In practice, employing parcellations as a data reduction technique prior to machine learning serves to aid in interpretability, reduces computational expense, and potentially provides regularizing benefits (i.e., fewer downstream features as considered in a classic understanding of the bias-variance trade-off, Von Luxburg 2011). That said, while brain parcellation techniques have proven useful and are widely popular, they suffer from a key limitation in contrast to finer spatial resolution analysis, namely that potentially useful information is lost when averaging across the vertices of an ROI.

A number of different parcellation schemes have been proposed, grouped roughly into structurally informed parcellations and functionally defined parcellations. Popular anatomically informed parcellations include the Desikan atlas, consisting of 34 bilateral cortical ROIs, and the Destrieux atlas, consisting of 74 cortical regions per hemisphere (Desikan 2006, Destrieux 2010). Functionally defined parcellations, emerging most prominently from the literature on resting state fMRI, tend to be data driven. Popular examples include the earlier Gordon atlas (Gordon et al., 2016), with parcel boundaries determined from resting state functional connectivity boundary maps, as well as the newer Schaefer parcellations, which further consider global similarity between parcels in addition to local boundary information (Schaefer, 2018).

### Choice of Parcellation for Machine Learning

The choice of parcellation or alternative dimensionality reduction strategy is often overlooked as a key parameter in building brain based-classifiers, or is undertaken in an *ad hoc* manner with a selection based on convenience or other unstated reason (Franke 2010, de Wit 2017, Squarcina 2017, Karch 2019, Gowin 2019, He 2020). Other parameters such as the choice of ML pipeline, feature selection or cross-validation strategy are often given more importance in exploring different experimental configurations (Mateos-Pérez 2018, Janssen 2018, Jollans 2019, Nielsen 2019). However, some studies have examined the choice of parcellation in more depth. Arslan et al. provide a systematic comparison across a number of surface-based parcellations and data reduction strategies as applied to resting state fMRI, determining that there was not one “best” parcellation (Arslan, 2018). With specific regards to downstream performance, Dadi et al. performed a thorough benchmarking with connectome-based predictions from fMRI (Dadi, 2019). This work identified functionally defined regions, as identified with data-driven clustering methods like dictionary learning, as the highest performing. Previous work on sMRI showed that increasing the spatial resolution of the cortical surface achieved higher performance when predicting age (Khundrakpam, 2015). The relationship between spatial scale and performance is one that we examine in more depth within this work.

### Research Goals

Here we investigated prediction of phenotypes from sMRI-derived measures projected onto the cortical surface under different summary parcellations. We investigated how choice of parcellation can influence predictive performance as well as interact with other modelling choices (e.g., does choice of “best” parcellation change under different ML strategies or for different predictive targets?). We also considered if there is an overall best strategy or combination of parcellations to obtain reliably high performance and what features make a parcellation predictive. We explored in particular the potential relationship between number of parcels and predictive performance, noting a potential power law scaling relationship. Likewise, we were interested in identifying the range of sizes over which a scaling relationship between size and performance might hold.

We also sought to characterize the potential gains in performance from employing strategies that can make use of information from multiple parcellations in order to inform predictions, such as ensembling. Specific questions we sought to address included which multiple parcellation strategy, as well as which parcellations are included in that strategy, and how those choices influence performance. For example, how do the number of parcellations as well as the number of parcels in each parcellation contribute to performance gains. Should the included parcellations for any one ensemble be all of one fixed size or instead span across different sizes (e.g., five parcellations of size 300 each versus five parcellations with sizes 100, 200, 300, 400 and 500). Finally, how these different decisions influence trade-offs between performance, runtime and interpretability, is an important consideration.

The core analysis of the paper was a systematic evaluation of different combinations of parcellations and ML pipelines. Each combination was scored based on its predictive performance across 45 different phenotypic variables. We conducted this investigation using data from the Adolescent Brain Cognitive Development (ABCD) Study, which provided sMRI surface data from 9,432 participants. Further, we evaluated how information across multiple parcellations can be leveraged to improve predictive performance, using both hyper-parameter and ensemble approaches.

## Methods

### Imaging Data

We used data from the ABCD Study^®^ release 2.0.1. Imaging data were sourced from the NDA Collection 3165 (See: https://collection3165.readthedocs.io/en/stable/). Data used within this study were the sMRI cortical reconstruction outputs of a modified HCP-style pipeline. For each participant, we used left and right hemisphere curvature, sulcal depth, cortical thickness and unsmoothed myelin map, each in the standard FS LR 32k vertex space (Glasser, 2013). We also employed each participant’s automatically computed FreeSurfer ROI-level summary statistics.

Initially, the subset of participants with available data across all four modalities of interest (e.g., curvature, sulcal depth, cortical thickness, and myelination) was determined, dropping any participant with missing data. Additional participant exclusions were applied based on a data driven outlier removal process, resulting in 9432 final participants. Exact details on outlier removal are available at sahahn.github.io/parc_scaling/outliers.

### Target Variables

A collection of 45 target phenotypic variables (23 binary and 22 continuous), used to gauge predictive performance of the different parcellation schemes, was sourced from release 2 of the ABCD dataset. Variables were sourced directly from the rds file made available by the Data Analysis, Informatics & Resource Center associated with the ABCD Study^®^. All collected variables, both target and brain, are from the baseline time point of the ABCD study^®^. Best efforts were made to source a list of representative and diverse variables. To facilitate the comparison of different parcellations, a larger list of variables, sourced from Owens 2021, was originally screened on a subset of the data (n=2000) to identify those showing some relationship with the sMRI measures as based on 5-fold ridge regression performance (see sahahn.github.io/parc_scaling/variables#is-predictive). As different target variables had highly non-overlapping amounts of missing data, we did not drop any participants and instead replaced all missing values with a missingness indicator. During evaluation, any target variables with missing values were excluded from both training and validation folds, and therefore did not contribute to the estimates of performance. As a result, the sample sizes available for each target variable varied slightly.

### Parcellations

All considered surface parcellations were in standard FS LR 32k space, to match the input data, and if necessary were re-sampled from their original space. We evaluated a number of existing surface parcellations and a number of randomly generated parcellations (details below). We also tested some additional variants including downsampled and FreeSurfer extracted region-of-interest values. In total, we assessed 82 existing parcellations. Parcellations available at multiple scales were assessed at every scale; in some cases where multiple versions of the same parcellation were available (e.g., from different resampling procedures or with different post processing applied), all versions were tested. 68 of the 82 parcellations were static or “hard” parcellations, in which each vertex is labelled as a part of exactly one parcel. We additionally considered 14 probabilistic or “soft” parcellations, where each parcel is represented by a set of probabilities or weightings across the whole surface or volume. As some parcellations were only originally available in volumetric MNI space, we applied registration fusion to map these parcellations to fsaverage space based on scripts available from Wu et al. (Wu, 2018). From fsaverage surface space, resampling to FS LR 32k space was conducted with tools available from the Human Connectome Project Workbench (Marcus, 2011). Depending on the type and original space of the parcellation a number of different strategies were necessary, and a more complete description of each option is available at sahahn.github.io/parc_scaling#resample_parcellations.

The existing parcellations were sourced from a number of different online repositories and in many cases have characteristics that may not be optimal for the present analyses. For example, some parcellations may not have been originally intended for sMRI (e.g., some were designed for resting state fMRI), some parcellations may have been created in a different standard space and therefore had to be re-sampled to the space required for the present analyses, and some parcellations may not have been originally intended to produce static parcellations. In order to partially address these concerns, we have included multiple versions of the same parcellation when different versions were available. That said, these limitations reflect the real-world challenges and options available currently to researchers.

We also generated random parcellations across a range of different scales. Random parcellations are generated as follows: For a random parcellation of size N, N locations were first selected at random across each hemisphere’s 59,412 vertices (with FS LR 32k space medial wall vertices automatically set to no parcel). Each location became the seed for a new contiguous region and was randomly assigned a size probability between 0 and 1, which is considered when choosing which region to grow. Next, a region was randomly selected according to those probabilities (i.e., a region assigned an initial probability of .5 would be picked on average twice as often as a region assigned .25). Then, one of the adjacent vertices to the region was selected at random and added to that region. This sequence, selecting a region and adding one vertex, was repeated until all vertices were assigned to a region.

We also tested 6 different downsampled icosahedron parcellations. These spanned in size from 42 to 1442 regions per hemisphere. Finally, we assessed the Desikan and Destrieux ROI values as extracted by FreeSurfer. These differ from the other tested parcellations both in how values are generated (FreeSurfer extracts values in an individual’s native space whereas we extract values from data warped to a common space) in addition to the surface modalities used (only average thickness, surface area and mean curvature were employed, which differed from the features used in the base analyses).

### Machine Learning Pipelines

We employed three base ML pipelines, each with classifier and regressor variants, as a representative sample of different popular and predictive ML strategies. All machine learning analyses were conducted with the Brain Predictability toolbox: a python based library designed to facilitate neuroimaging based ML (Hahn, 2020). The first step in each pipeline was a loading component responsible for extracting data for the ROIs of the specified surface parcellation. The output from this step was a vector of concatenated values for each of the surface metrics (curvature, sulcal depth, thickness and myelin measures) in each ROI, generating a feature vector of length four times the number of parcels for each participant. Next, the ROI values were scaled using robust scaling, wherein each feature was standardized by subtracting the median and then scaling according to the 5th and 95th percentiles of that feature’s distribution. These features were then used as input to train a classifier or regressor under one of three different base configurations, which included:

#### Elastic-Net

The base ML estimator within the pipeline under this configuration was a logistic or linear regression with elastic-net penalty available from scikit-learn (Pedregosa, 2011). A nested random hyper-parameter search (Random Search) over 60 combinations was evaluated through a nested 3-fold CV to select the strength of regularization applied as well as the ratio between l1 and l2 regularization.

#### SVM

The base ML estimator within the pipeline under this configuration was a Support Vector Machine (SVM) classifier or regressor with radial basis function kernel available from scikit-learn (Amari 1999). A front-end univariate feature selection procedure was added to this pipeline (based on an ANOVA F-value between a feature and the target variable, keeping only a percentage of the top features). A nested random hyper-parameter search (Random Search) over 60 combinations was then evaluated through a nested 3-fold CV in order to select the SVM’s strength of regularization and kernel coefficient as well as the percentage of features to keep in the front-end feature selector. All three hyper-parameters were optimized simultaneously. Notably, the feature selection step could remove no more 50% and no fewer than 1% of the passed features.

#### LGBM

The base ML estimator was an extreme gradient boosted tree-based classifier and regressor from the Light Gradient Boosting Machine (LGBM) package (Chen, 2015). The tuned hyper-parameters for this algorithm included the type of boosting, the number of estimators, different tree sampling parameters, and regularization parameters. Given the high number of hyper-parameters to tune in contrast to the other base estimators (nine), we employed a two-point differential evolution-based hyper-parameter search strategy (Two Points DE) implemented through the python library Nevergrad (Rapin 2018). The search was run for 180 iterations, in which each set of parameters was evaluated with a single 25% nested validation split.

More detailed information on ML pipeline implementation can be found at sahahn.github.io/parc_scaling/ml_pipelines.

### Evaluation

To evaluate a given target variable, parcellation, or machine learning strategy’s performance, we defined an explicit framework to compare different combinations of methods. We evaluated each combination of target variable, parcellation and ML pipeline with five-fold cross-validation using the full set of available participants (Figure 1). Each of the validation folds, including any nested parameter tuning folds, were conducted such that participants from the same family were preserved within the same training or testing fold. The 5-fold structure was kept constant and therefore comparable across all combinations of ML pipeline, target variable, and parcellation. In the case of missing target variables, those participants with missing data were simply excluded from their respective training or validation fold. This strategy was used for each of the proposed combinations to generate metrics of performance: explained variance for regression predictors and area under the receiver operator characteristic curve (ROC AUC) for binary predictors.

**Figure 1:**
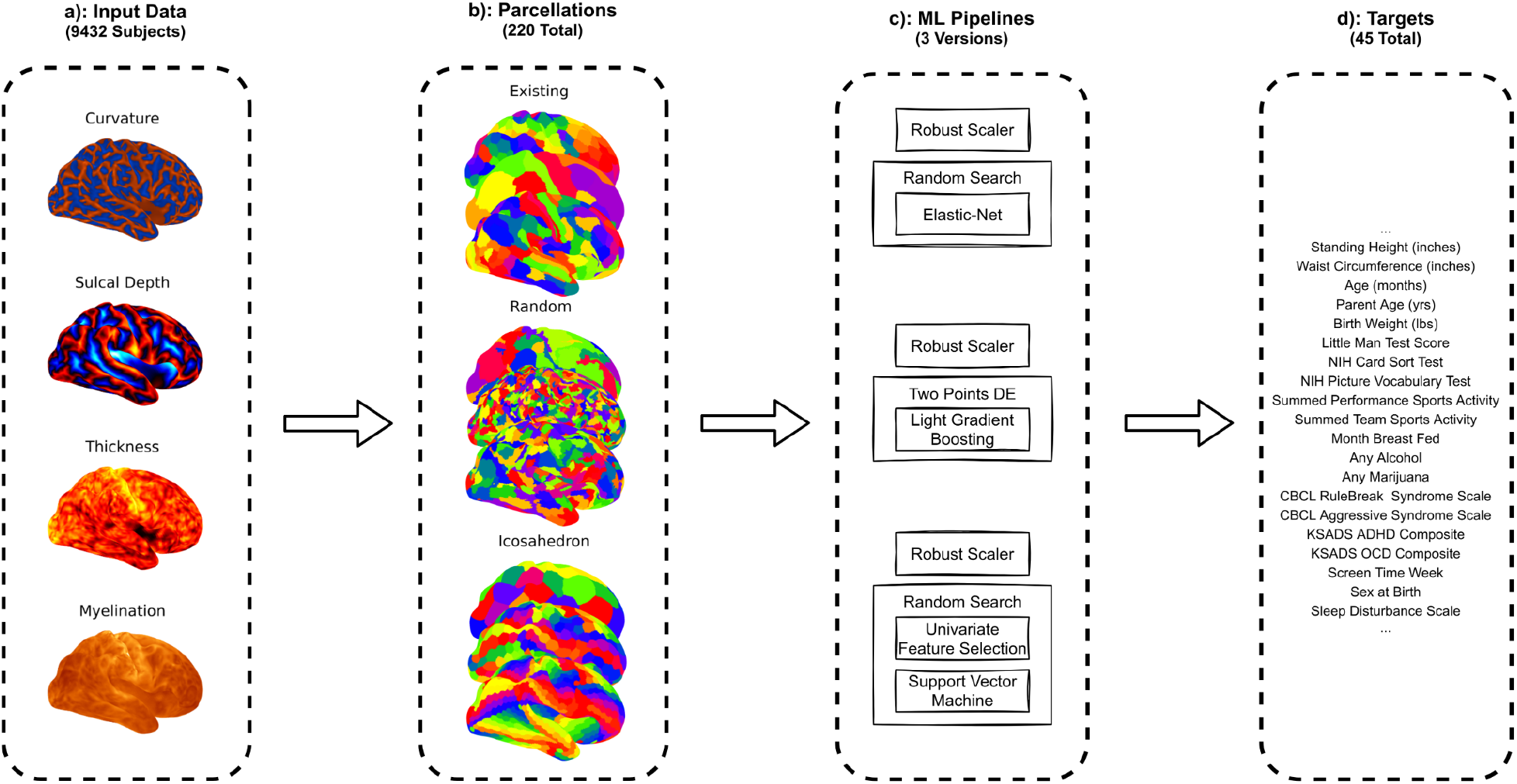
Diagram outlining the core analytic procedure conducted where combination of parcellation, ML pipeline and target variable were systematically evaluated. Section a. shows the four different sMRI measures used as input data for each participant. Section b. highlights that each of the four measures were parcellated into mean region-of-interest values for 220 different parcellations. Section c. shows the three different ML pipelines, each based on a different base ML estimator, which were tested on each parcellation-target pair. Section d. lists a subset of the 45 different target variables for which out-of-sample predictive utility was estimated for each parcellation-pipeline pair.

### Multiple Parcellation Strategies

In addition to the base analysis described above, we sought to quantify how additional strategies operating across multiple parcellations might perform. Considering multiple parcellations at once has the benefit of potentially capturing overlapping information present across different spatial scales, and could therefore prove superior to adhering to just one scheme. Given a potentially limitless number of potential configurations, we explored only a small subset. These extensions to the base analysis can be separated into three different analysis types: choice of parcellation as a nested hyper-parameter, ensembling over multiple parcellations using voting, and ensembling using stacking (Wolpert, 1992). In all of these analytic approaches, randomly generated parcellations were used as a virtually limitless number can be generated at any desired spatial scale.

In order to treat choice of parcellation as a hyperparameter, we employed a nested grid search. A three-fold nested cross-validation scheme on the training set, respecting family structure as before (i.e., assigning members of the same family to the same fold), was used to evaluate each potential parcellation. Within each of these internal folds a ML pipeline was trained, with its own nested parameter tuning, and then evaluated on its respective internal validation set. This process yielded an average for each of the three folds’ scores for each parcellation. The parcellation which obtained the highest score was selected for re-training on the full training set which involved, as in each nested fold, training a ML pipeline with its own nested parameter search. The final trained ML estimator, with the selected best parcellation, was then evaluated on the validation fold. This process was repeated across the whole training set according to the same five-fold structure as used in the base analyses, thus allowing the results to be directly comparable.

In ensemble analyses, we tested two different ensemble strategies: voting and stacking. In the voting ensemble approach, a separate ML pipeline was trained for each available parcellation, where each individual pipeline-parcellation pair was trained in the same way as in the base analysis. To do this, each trained ML pipeline from the previous step first generated a prediction. Then, the voting ensemble aggregated the predictions as either the mean, in the case of regression, or the most frequently predicted class, in the case of classification. The aggregated scores were then scored as a single set of predictions.

The stacking ensemble, while similar to the voting ensemble, is more complex. For each of the pipeline-parcellation combinations, a separate three-fold cross-validation framework was used in the training set. In this framework, three ML pipelines were trained on 2/3 of the training set and predictions were made for the remaining 1/3, ultimately yielding an out-of-sample prediction for each participant in the training set. The predictions from all pipeline-parcellation combinations were used as features to train a “stacking model”. The purpose of the stacking model was to learn a relative weighting of each parcellation-pipeline combination (i.e., to give more weight to “better” parcellation-pipeline combinations and less weight to “worse” ones). The algorithm used to train the stacking model was a ridge penalized linear or logistic regression with nested hyper-parameter tuning. Once trained, this stacking model was used to predict the target variable in a novel sample (i.e., the held-out test set). The stacking ensemble procedure notably involved a large increase in computation relative to the voting ensemble, as the stacking ensemble involved training three pipelines for each parcellation-pipeline combination, whereas the voting ensemble consisted of training only one ML pipeline for each.

For the number of different random parcellations available to a search or ensemble strategy, we evaluated four different numbers of parcellations: 3, 5, 8 and 10. Further, for each of these numbers of parcellations, we tested fixed size parcellations as well as differentially sized parcellations across a range of sizes (100, 200, 300, 400, 500, 50-500, 100-1000 and 300-1200). For example, for a combination of 3 parcellations and a fixed size of 100, three random parcellations with size 100 could be used. For a combination of 5 parcellations of a range of sizes from 100-1000, parcellations of size 100, 325, 550, 775 and 1000 could be used. All combinations are then repeated twice with two different versions of parcellations at each size used.

In summary, all possible combinations of the following parameters were evaluated:

- 8 Size Configurations (5 Fixed Sizes + 3 Across Sizes)
- 4 Numbers of Parcellations (3, 5, 8, 10)
- 3 Base Strategies (Grid, Voted, Stacked)
- 4 Pipelines (3 Base Pipelines + ‘All’ Configuration)
- 45 Target Variables
- 2 Random Repeats

### Multiple Parcellation Evaluation

In the base analysis, each parcellation was evaluated with five-fold cross-validation across all combinations of target and ML pipeline. Here, we evaluate each additional multiple parcellation strategy, Grid (choice of parcellation as a hyper-parameter), Voted (voting ensemble) and Stacked (stacking ensemble) with the same five-fold cross-validation across the same combinations of target and ML pipeline. This setup allowed us to compare directly between using a single fixed parcellation versus information across multiple parcellations. Additionally, we also considered a special ‘All’ configuration, ensembling and selecting across both parcellation and choice of ML pipeline (e.g., a voting ensemble which averages predictions from SVM, Elastic-Net and LGBM pipelines, each trained on random parcellations of size 100, 200 and 300). This configuration provides the potential to exploit information from not just multiple parcellations, but also the unique information generated from each ML pipeline.

### Mean Rank

In order to evaluate and compare across the different binary and regression metrics, as well as to address scaling issues between metrics (e.g., sex is more predictable than the ADHD composite score), we adopted mean rank as our ultimate performance metric of interest. We computed the relative per-target ranking across parcellations (or in some cases parcellation-pipeline combinations), where the parcellation result with the highest score would receive rank 1 (i.e., lower rank better). This metric is therefore sensitive to the collection of included parcellations or parcellation-pipeline pairs. Mean rank can also refer to different aggregations of these base ranks. For example in Figures 2 and 5 an average rank is first computed separately for each pipeline across all the targets, then set as a final mean rank across pipeline averages. In contrast, for example in the bottom of Figure 3, mean rank is computed directly between pairs of parcellation-pipelines.

**Figure 2:**
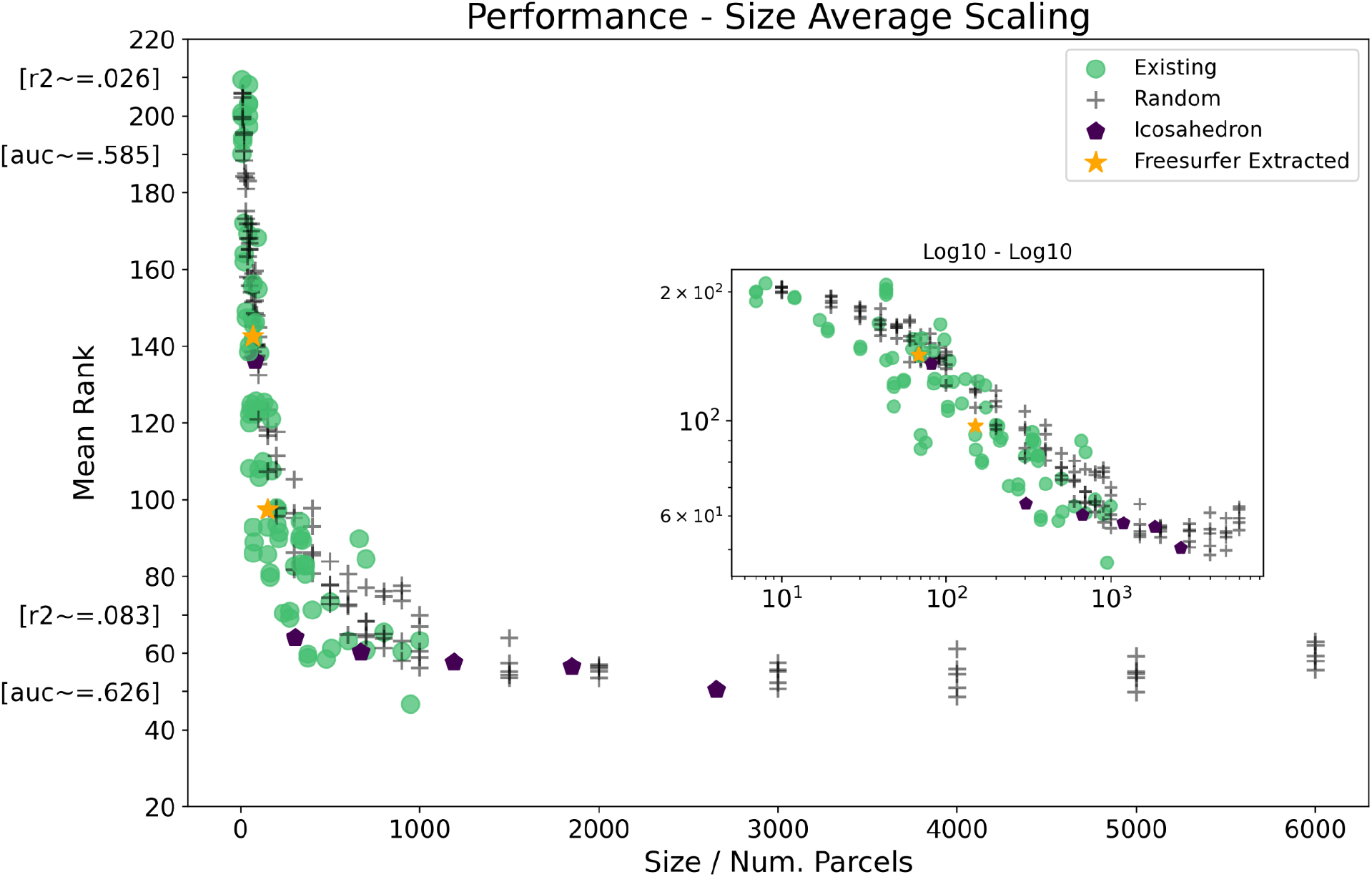
Performance represented by mean relative ranking between 220 parcellations, against the number of parcels in each parcellation. Shape and color of each point indicates the type of parcellation. The inset figure shows the main figure on a log10 scale. Green circles indicate that a parcellation is from an existing source, either static or probabilistic. Gray plus markers indicate that the parcellation was randomly generated. Purple pentagons indicate that the parcellation represents a down-sampled icosahedron. The two orange stars indicate the parcellation values as extracted directly from Freesurfer. Example estimates of R2 and ROC AUC are also provided on the y-axis as estimated from the means obtained across the 22 regression or 23 binary target variables of as close as possible to that rank.

**Figure 3.**
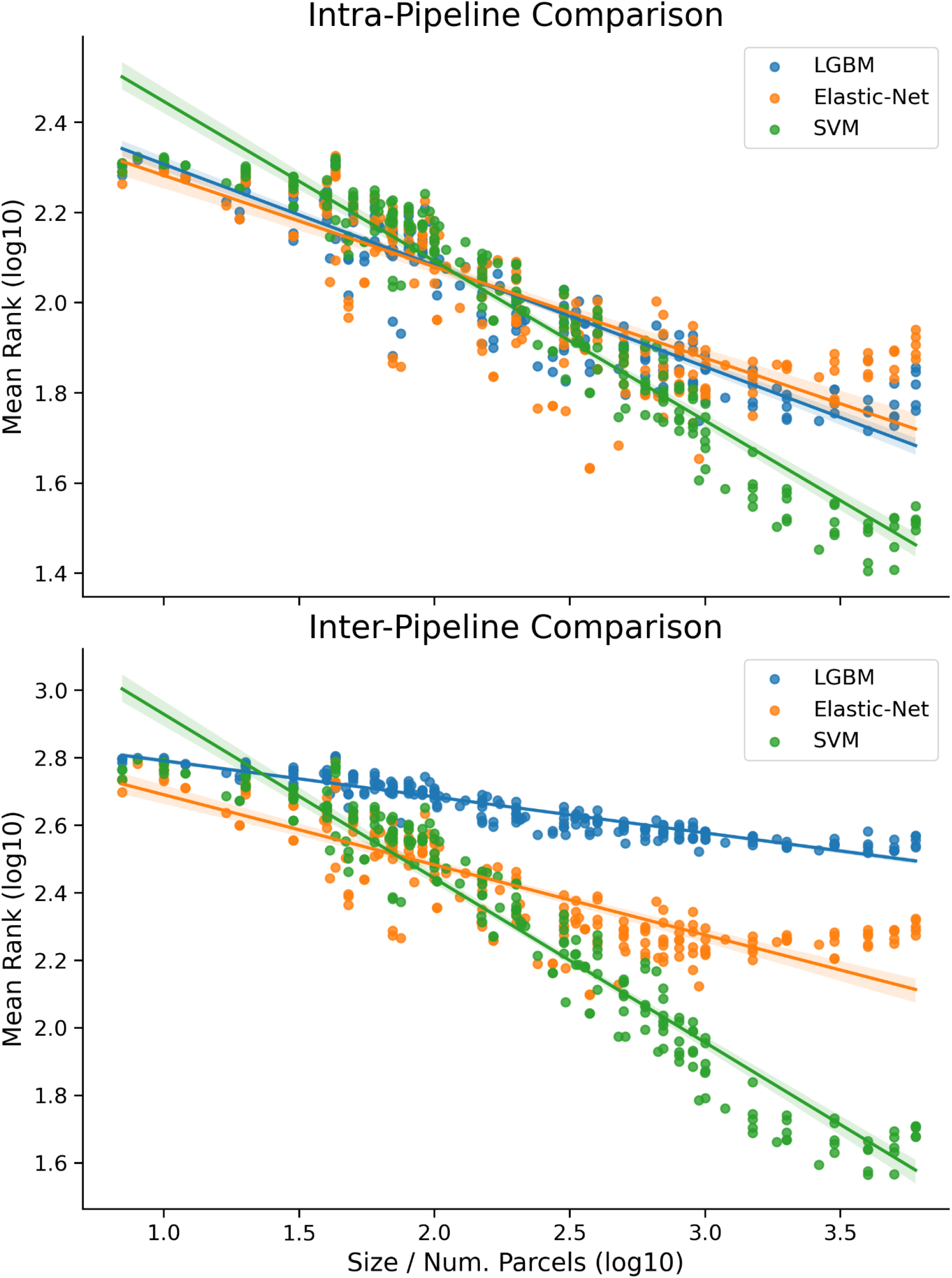
shows the log10-log10 relationship between number of parcels and mean rank calculated in two different ways. The top figure, Intra-Pipeline Comparison, shows mean rank for each pipeline computed relative only to other parcellations evaluated with the same pipeline. Comparisons were made between 220 (i.e., maximum mean rank is 220 or on log10 y-axis 2.34) parcellations for each pipeline. In contrast, the bottom figure, Inter-Pipeline Comparison, shows the mean rank calculated between each parcellation-pipeline combination. Comparisons were made between 660 (2.82 on log10 y-axis) parcellation-pipeline combinations. Each parcellation-pipeline result is colored by its underlying pipeline type, regardless of type of parcellation. Robust regression lines of best fit on the log10-log10 data are plotted separately for each pipeline across both figures (shaded regions around the lines of fit represent the bootstrap estimated 95% CI).

### Modelling Results

We employ ordinary least squares regression (OLS), as implemented in the python package statsmodel (Seabold 2010), to model results from the base experiments. Base notation for OLS equations are written as *A* ∼ *B* + *C* where A is the dependent variable and B + C are independent fixed effects. Alternatively if written as *A* ∼ *B* * *D* then D will be added as a fixed effect along with an interaction term between B and D (equivalent to alternative notation *A* ∼ *B* + *D* + *B* * *D*). If a fixed effect was categorical, then it was dummy coded and each dummy variable added as a fixed effect. Last, if a variable was within the brackets of log10(), then the logarithm of the variable with base 10 was used. For example, if the dependent variable was Mean Rank with fixed effects of the log of Parcellation Size and which ML Pipeline was used, we would write *Mean Rank* ∼ *log*10(*Parcellation Size*) + *Pipeline*.

### Computations

The computations for this work were performed on the Vermont Advanced Computing Core, a high performance computing cluster made available to researchers. All code and shareable data are available at sahahn.github.io/parc_scaling which further includes instructions on how to download any non-sharable data.

## Results

### Averaged Performance-Size Scaling

The results from the core analysis with a focus on the relationship between performance and number of parcels in each parcellation scheme are shown in Figure 2. Specifically, we consider here the mean rankings as averaged across all different base ML pipelines as well as target variables. Notably, the best (lowest) mean rank observed was the downsampled version of the probabilistic DiFuMo 1024 scale parcellation, with mean rank 46.88 (Dadi 2020). The two Freesurfer extracted parcellations obtained mean ranks of Desikan=142.59 and Destrieux=97.42.

We first estimated the range of parcellation sizes where a power-law like distribution best fit the data to be 4000 parcels and fewer (a description of this procedure is described at https://sahahn.github.io/parc_scaling/estimate_powerlaw.html, Clauset, 2007). In order to model the relationship between parcellation size and performance while accounting for type of parcellation we fit an OLS model as, *log*10(*mean rank*) ∼ *log*10(*size*) + *parcellation type*, on parcellations with 4,000 or fewer parcels. Model fit statistics are reported in Table 3, where notably the overall model fit was *R*^*2*^=0.90. Of further interest, the estimated coefficient for parcellation size (i.e., the scaling exponent in a power law distribution) was 0.-2753 and highly significant. The only significant effect for parcellation type was if a parcellation was randomly generated (*coef*=.0488). No other significant effects between subgroups of parcellations were found, although this may be due to the small number of samples in these subgroups (e.g., only 6 icosahedron parcellations). Further, a separate OLS allowing for interactions between the number of parcels and parcellation type did not reveal any significant interactions.

**Table 1.** lists all the target variables employed in these analyses. The first half of the table lists the 22 continuous variables and the second half lists the 23 binary variables, with the latter further broken down by class value. The participants excluded for missing data refers to the number of missing values for that variable, which are not included in any of the provided summary statistics, and those participants are likewise skipped when evaluating that target variable. Full descriptions and extra information on each target variable are included with the online supplementary materials (sahahn.github.io/parc_scaling/variables).

**Participants excluded for missing data**
|  |  |  |  |
| --- | --- | --- | --- |
| Standing Height (inches) | 9427 | 55.27 ± 3.32 | 5 |
| Waist Circumference (inches) | 9420 | 26.51 ± 4.74 | 12 |
| Measured Weight (lbs) | 9425 | 82.18 ± 23.24 | 7 |
| CBCL RuleBreak Syndrome Scale | 9425 | 1.16 ± 1.82 | 7 |
| Parent Age (yrs) | 9364 | 40.08 ± 6.72 | 68 |
| Motor Development | 9340 | 2.37 ± 0.8 | 92 |
| Birth Weight (lbs) | 9032 | 6.57 ± 1.48 | 400 |
| Age (months) | 9432 | 119.17 ± 7.45 | 0 |
| Little Man Test Score | 9163 | 0.59 ± 0.17 | 269 |
| MACVS Religion Subscale | 9429 | 3.32 ± 1.42 | 3 |
| Neighborhood Safety | 9426 | 3.93 ± 0.95 | 6 |
| NeuroCog PCA1 (general ability) | 8791 | 0.04 ± 0.76 | 641 |
| NeuroCog PCA2 (executive function) | 8791 | 0.02 ± 0.76 | 641 |
| NeuroCog PCA3 (learning / memory) | 8791 | 0.03 ± 0.7 | 641 |
| NIH Card Sort Test | 9312 | 92.89 ± 9.27 | 120 |
| NIH List Sorting Working Memory Test | 9282 | 97.15 ± 11.72 | 150 |
| NIH Comparison Processing Speed Test | 9297 | 88.4 ± 14.48 | 135 |
| NIH Picture Vocabulary Test | 9315 | 84.88 ± 7.93 | 117 |
| NIH Oral Reading Recognition Test | 9305 | 91.08 ± 6.72 | 127 |
| WISC Matrix Reasoning Score | 9234 | 18.03 ± 3.77 | 198 |
| Summed Performance Sports Activity | 9432 | 0.99 ± 1.03 | 0 |
| Summed Team Sports Activity | 9432 | 1.17 ± 1.18 | 0 |

| <u>Binary Variables</u> | <u>Participants</u><br><u>included</u> | <u>Freq.</u> | <u>Participants</u><br><u>excluded for</u><br><u>missing data</u> |
| --- | --- | --- | --- |
| Speaks Non-English Language | 9429 | 1.0 | 3 |
| No | 6399 | 0.68 | - |
| Yes | 3030 | 0.32 | - |
| Thought Problems ASR Syndrome Scale | 9432 | 1.0 | 0 |
| ≤2 | 7585 | 0.8 | - |
| >2 | 1847 | 0.2 | - |
| CBCL Aggressive Syndrome Scale | 9432 | 1.0 | 0 |
| ≤4 | 7032 | 0.75 | - |
| >4 | 2400 | 0.25 | - |
| Born Premature | 9333 | 1.0 | 99 |
| No | 7541 | 0.81 | - |
| Yes | 1792 | 0.19 | - |
| Incubator Days | 9432 | 1.0 | 0 |
| 0 | 8203 | 0.87 | - |
| >0 | 1229 | 0.13 | - |
| Months Breast Fed | 9432 | 1.0 | 0 |
| ≤10 | 6497 | 0.69 | - |
| >10 | 2935 | 0.31 | - |
| Has Twin | 9401 | 1.0 | 31 |
| No | 7507 | 0.8 | - |
| Yes | 1894 | 0.2 | - |
| Planned Pregnancy | 9236 | 1.0 | 196 |
| No | 3511 | 0.38 | - |
| Yes | 5725 | 0.62 | - |
| Distress At Birth | 9432 | 1.0 | 0 |
| 0 | 6853 | 0.73 | - |
| >0 | 2579 | 0.27 | - |
| Mother Pregnancy Problems | 9432 | 1.0 | 0 |
| 0 | 5492 | 0.58 | - |
| >0 | 3940 | 0.42 | - |
| Any Alcohol During Pregnancy | 9432 | 1.0 | 0 |
| No | 7470 | 0.79 | - |
| Yes | 1962 | 0.21 | - |
| Any Marijuana During Pregnancy | 9432 | 1.0 | 0 |
| No | 9102 | 0.97 | - |
| Yes | 330 | 0.03 | - |
| KSADS OCD Composite | 9432 | 1.0 | 0 |
| 0 | 8457 | 0.9 | - |
| >0 | 975 | 0.1 | - |
| KSADS ADHD Composite | 9432 | 1.0 | 0 |
| 0 | 7486 | 0.79 | - |
| >0 | 1946 | 0.21 | - |
| KSADS Bipolar Composite | 9432 | 1.0 | 0 |
| 0 | 8056 | 0.85 | - |
| >0 | 1376 | 0.15 | - |
| Mental Health Services | 9376 | 1.0 | 56 |
| No | 7923 | 0.85 | - |
| Yes | 1453 | 0.15 | - |
| Detentions / Suspensions | 9248 | 1.0 | 184 |
| No | 8786 | 0.95 | - |
| Yes | 462 | 0.05 | - |
| Parents Married | 9361 | 1.0 | 71 |
| No | 2856 | 0.31 | - |
| Yes | 6505 | 0.69 | - |
| Prodromal Psychosis Score | 9432 | 1.0 | 0 |
| <=10 | 7689 | 0.82 | - |
| >10 | 1743 | 0.18 | - |
| Screen Time Week (hrs) | 9432 | 1.0 | 0 |
| <=4 | 8265 | 0.88 | - |
| >4 | 1167 | 0.12 | - |
| Screen Time Weekend (hrs) | 9432 | 1.0 | 0 |
| <=5 | 7430 | 0.79 | - |
| >5 | 2002 | 0.21 | - |
| Sex at Birth | 9427 | 1.0 | 5 |
| Female | 4532 | 0.48 | - |
| Male | 4895 | 0.52 | - |
| Sleep Disturbance Scale | 9432 | 1.0 | 0 |
| <=35 | 5295 | 0.56 | - |
| >35 | 4137 | 0.44 | - |

**Table 2.** This table shows the source for all collected “existing” parcellations. Included for each parcellation in the far left column is the corresponding name or shortened name of the parcellation. The second column lists the number of sourced parcellations that fall under this name to be used, where numbers greater than one refer to either different versions of the same parcellation, or parcellations spanning multiple spatial resolutions. The third column lists if the parcellations considered are ‘Hard’ (i.e., every vertex is assigned to a parcel) or ‘Soft’ (i.e., parcels are represented by probabilistic maps). Last, the fourth column lists the reference to where the parcellation was sourced if available.

| **<u>Name</u>** | **<u># of Parcellations</u>** | **<u>Type</u>** | **<u>Reference</u>** |
| --- | --- | --- | --- |
| Schaefer Local-global parcellation | 10 (scales 100-1000) | Hard | Schaefer 2018 |
| Gordon | 3 (different sources) | Hard | Gordon 2016 |
| Brodmann Areas | 1 | Hard | Brodmann 1909 |
| VDG11b | 1 | Hard | Van Essen 2012 |
| HCP-MMP | 3 (different sources) | Hard | Glasser 2016 |
| Automatic Anatomical Labeling (AAL) | 2 (different sources) | Hard | Tzourio-Mazoyer 2002 |
| Baldassano | 1 | Hard | Baldassano 2015 |
| Desikan | 2 (different sources) | Hard | Desikan 2006 |
| Destrieux | 2 (different sources) | Hard | Destrieux 2010 |
| Brainnetome | 2 (different sources) | Hard | Fan 2016 |
| Power | 2 (different sources) | Hard | Power 2011 |
| Shen 268 Parcels | 2 (different sources) | Hard | Shen 2013 |
| Shen 368 Parcels | 1 | Hard | Salehi 2020 |
| Yeo | 3 (7 Networks, 17 Networks and parcel level version) | Hard | Yeo 2014 |
| DiFuMo | 5 (scales 64-1028) | Soft | Dadi 2020 |
| MIST | 9 (scales 9-444) | Hard | Urchs 2019 |
| AICHA | 1 | Hard | Joliot 2015 |
| Economo | 1 | Hard | von Economo 2915 |
| NSPN500 | 1 | Hard | Whitaker 2016 |
| Oasis | 1 | Hard | Sabuncu 2011 |
| SJH | 1 | Hard | Harrison 2015 |
| Allen | 1 | Soft | Allen 2011 |
| BASC | 9 (scales 9-444) | Hard | Bellec 2013 |
| MSDL | 1 | Soft | Varoquaux 2011 |
| Harvard Oxford | 4 (different versions) | Hard / Soft | Jenkinson 2012 |
| Craddock | 4 (different versions) | Soft | Craddock 2012 |
| Smith ICA | 2 (different versions) | Soft | Smith 2009 |
| CPAC | 1 | Hard | Craddock 2013 |
| Hammersmith | 1 | Hard | Hammers 2003 |
| JuBrain | 1 | Hard | Eickhoff 2005 |
| MICCAI | 1 | Hard | http://www.neuromorphometrics.com/2012_MICCAI_Challenge_Data.html |
| Slab | 2 (907 and 1068) | Hard | Sripada 2014 |

**Table 3:**
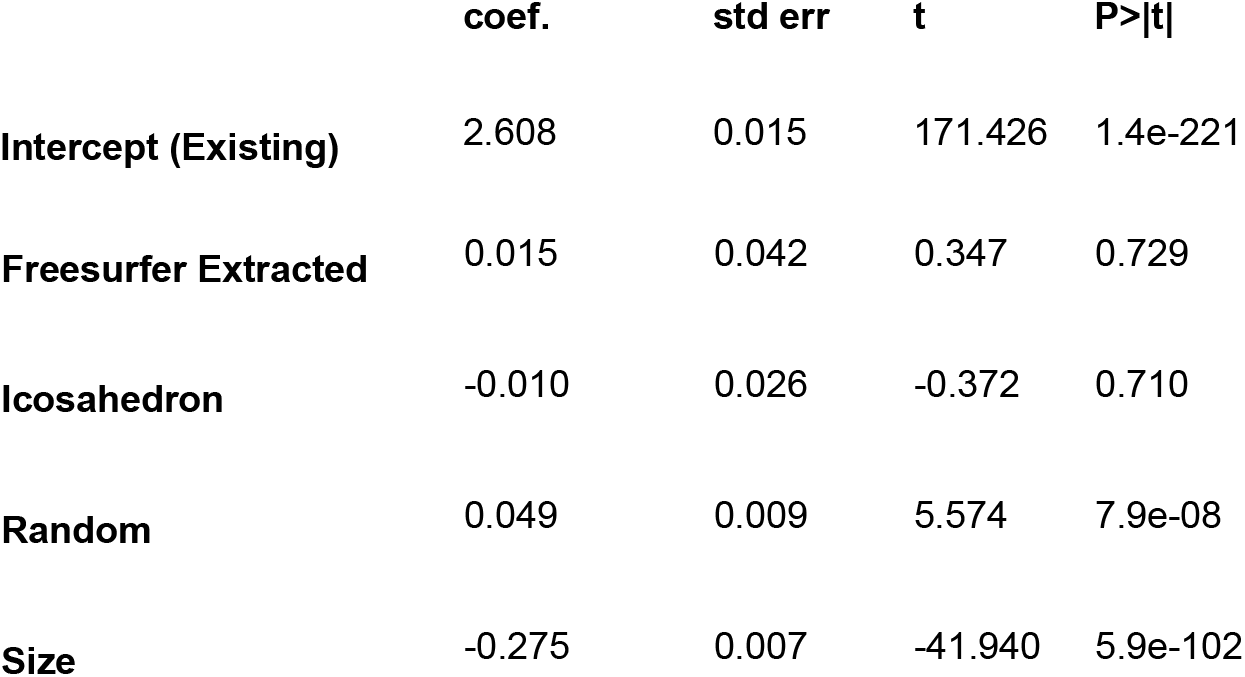
OLS model *log*10(*mean rank*) ∼ *log*10(*size*) + *parcellation type* fit on all parcellation results of size 4,000 and less, corresponding to the results plotted in Figure 2. As parcellation type was a categorical variable, it was dummy coded with Existing type parcellations as the intercept and separate model coefficients (coef.) generated for the additional parcellation types (Freesurfer, Icosahedron and Random), as well as for Size. Model fit *R*^*2*^=0.90, *F*=459.2, *P(F)*=1.79e-100.

|  | coef. | std err | t | P>\|t\| |
| --- | --- | --- | --- | --- |
| Intercept (Existing) | 2.608 | 0.015 | 171.426 | 1.4e-221 |
| **Freesurfer Extracted** | 0.015 | 0.042 | 0.347 | 0.729 |
| **Icosahedron** | -0.010 | 0.026 | -0.372 | 0.710 |
| **Random** | 0.049 | 0.009 | 5.574 | 7.9e-08 |
| **Size** | -0.275 | 0.007 | -41.940 | 5.9e-102 |

### Choice of Pipeline

The top panel of Figure 3 shows the relationship between observed performance to number of parcels changes when estimated using different ML pipelines. We estimated the region where power law scaling holds separately for each of the three pipelines in order to provide a more fair comparison between pipelines (LGBM: 7-4000, Elastic-Net: 7-1500, SVM: 20-4000). The OLS model (Table 4 - left) was then fit with only the results within these estimated ranges. The base estimated coefficient for the number of parcels (Size) was -.2651, similar to the previous estimate. The SVM pipeline significantly differed from the referent Elastic-Net with coefficient .3104, whereas the LGBM pipeline did not. These results indicate that there are differences between the pipelines (i.e., size/scaling coefficient, range of scaling and intercept), as well as confirm more generally that scaling, albeit with varying degree, holds regardless of pipeline.

**Table 4:** Both the left and right of the table present summary results from OLS model *log*10(*mean rank*) ∼ *log*10(*size*) * *pipeline*. The key difference is that on the left (Intra-Pipeline) mean rank is calculated within each pipeline separately and on the right (Inter-Pipeline) mean rank is calculated only once across all parcellation-pipeline combinations. The Intra-Pipeline (left) model fit was R^2^=.877, F=829.6, P(F)=2.74e-262. The Inter-Pipeline (right) model fit was R^2^=0.921, F=1522, P(F)<1.0e-300.

|  | Intra-Pipeline |  |  |  | Inter-Pipeline |  |  |  |
| --- | --- | --- | --- | --- | --- | --- | --- | --- |
|  | coef. | std err | t | P>\|t\| | coef. | std err | t | P>\|t\| |
| **Intercept (Elastic-Net)** | 2.598 | 0.020 | 133.247 | <1e-300 | 2.884 | 0.018 | 160.158 | <1e-300 |
| **LGBM** | -0.032 | 0.026 | -1.231 | 0.219 | 0.017 | 0.025 | 0.422 | 0.673 |
| **SVM** | 0.310 | 0.029 | 10.801 | 1.8e-25 | 0.520 | 0.025 | 20.432 | 3.9e-72 |
| **Size (Elastic-Net)** | -0.265 | 0.009 | -30.051 | 2.3e-122 | -0.202 | 0.007 | -27.342 | 2.4e-110 |
| **Size \* LGBM** | 0.022 | 0.012 | 1.923 | 0.055 | 0.097 | 0.010 | 9.267 | 2.7e-19 |
| **Size \* SVM** | -0.132 | 0.012 | -10.662 | 1.5e-25 | -0.280 | 0.010 | -26.707 | 8.0e-107 |

The bottom panel of Figure 3 and right side of Table 4 show another comparison between pipelines, but with an alternative definition of mean rank that allows for an inter-pipeline comparison. Rank here is now relative to all other pipeline-parcellation pairs (660 total), whereas before ranks were calculated separately for each pipeline. The base size coefficient (Elastic-Net slope) was -0.203, differing significantly in the interaction term for both SVM -0.2804 and LGBM 0.0973. The estimated intercept (Elastic-Net), 2.886, significantly differed only for the SVM pipeline (coef=0.521) and not the LGBM pipeline. A potential point of interest is the area around size 100 where the line of fit for the Elastic-Net and SVM based results intersect, marking a transition point where the SVM starts to outperform Elastic-Net based pipelines. We note that while internal scaling (i.e., that some relationship between performance and size holds despite choice of pipeline) may be mostly consistent, as shown in the top of Figure 3, when allowing for explicit comparisons between pipelines we find significant differences in both performance and the slope of size to performance.

### Variation Across Target Variable

Whereas the previous figures showed mean rank computed across all target variables, Figure 4 shows the extent to which the results varied across the 45 different target variables. Notably, we can compare the model fit of a simple OLS model, *log*10(*mean rank*) ∼ *log*10(*size*), between the target specific ranks (R2=0.482) and the mean across targets (R2=0.883). That is to say, the degree to which the performance gains are explained by parcellation size are more consistent on average, but case to case exhibit more variance. Visually, the observed variance, or rather deviance of target specific ranks from the mean, appears to increase at larger parcellation sizes. To formally characterize the general pattern of increasing spread in mean rank (across targets) as the number of parcels grows, we first calculated the interquartile range (IQR) of log10 mean rank at each unique parcellation size. Next, we fit an OLS model, *IQR* ∼ *log*10(*size*), with fit R2=0.796, F=312.3, P(F)=2.38e-29, and significant size coefficient (slope) of 0.176.

**Figure 4.**
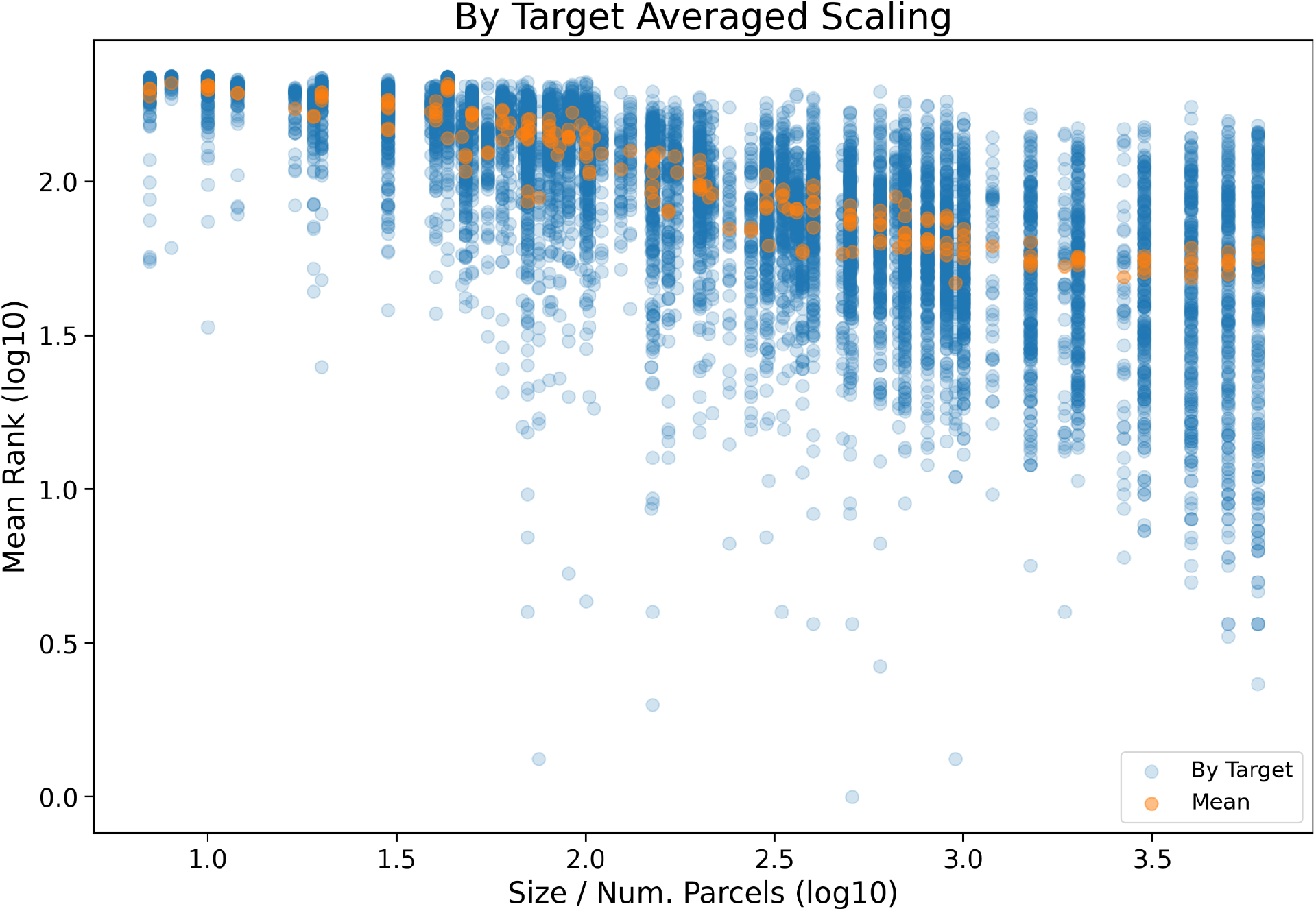
shows for each parcellation the computed log10 mean rank across pipeline for each of the 45 target variables (Blue), as well as the mean rank across all targets (Orange). Results are plotted on the x-axis according to their log10 number of parcels in the corresponding parcellation.

### Single vs. Multiple Parcellations

Figure 5 compares single parcellation results versus different types of multiple parcellation results. Size, as plotted on the x-axis, for ensemble based parcellations was calculated as the sum of base parcels used, which was done in order to capture the total number of unique parcellations contributing to that predictive model. Alternatively, the sizes for the ‘Grid’ hyper-parameter based results were set to match the largest size parcel considered as ultimately only a single parcellation contributes to the prediction. With the introduction of the multiple parcellation strategies, we can now test the following four ideas:

**Figure 5.**
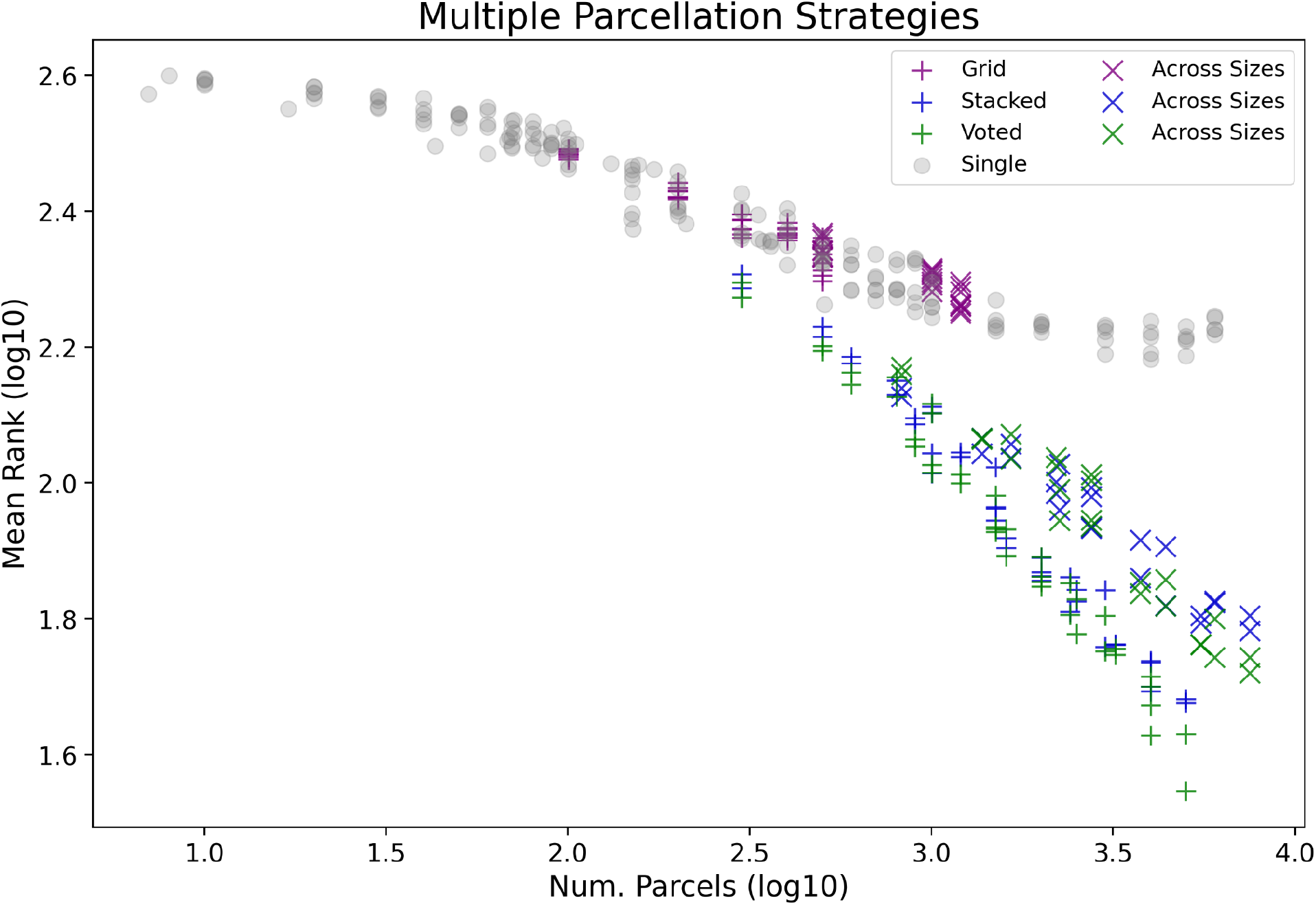
plots the log10 mean parcellations rank as averaged across pipelines separated by different parcellation strategies. This figure includes the single parcellation results from Figure 2 plotted as gray dots, as well as the three multiple parcellation strategies (Grid - pink, Voted - green and Stacked - blue). For the multiple parcellation strategies, results are further split into those in which the pool of parcellations were sourced from random parcellations of the same size (+), or sourced from a range of parcellations across different spatial scales (x)- Results are plotted on the x-axis according to their log10 number of parcels, where number of parcels for the ensemble based methods are calculated as a sum of the ensemble parcellations. Alternatively, for the grid search, results were plotted under the highest number of parcels that search had access to (e.g., if between sizes 100, 200 and 300, it would be plotted under size 300).

1. Are ensembles of parcellations better than non-ensembled strategies? The results from Table 5, specifically the interaction between size and if ensemble based, indicated that ensembles of parcellations were better than the single parcellation based results.
2. Are voting or stacking based ensembles better? An OLS model fit on only the subset of ensemble results found no significant differences between the voting and stacking based ensembles (p>|t|=0.320).
3. Are ensembles of fixed size random parcellations or random parcellations from a range of sizes better? A significant reduction in performance was observed when parcellations were sourced from across sizes vs. from fixed sizes (across sizes coef.=0.141, relative to fixed sizes intercept=3.276).
4. Does the ‘Grid’ hyper-parameter based multiple parcellation selection perform better than selecting a single random parcellation of similar size? An OLS model fit on a subset of just the ‘Grid’ and random parcellation results between sizes 100 and 1200 indicated that there were no significant differences.

**Table 5:** OLS model *log*10(*mean rank*) ∼ *log*10(*size*) + *is ensemble* fit on all results as plotted in Figure 5. Model fit *R*^*2*^=0.96, *F*=3211, *P(F)*=2.5e-283.

|  | coef. | std err | t | P>\|t\| |
| --- | --- | --- | --- | --- |
| **Intercept (Not Ensemble)** | 2.786 | 0.011 | 248.940 | <1e-300 |
| **Is Ensemble** | 0.615 | 0.046 | 13.348 | 5.6e-34 |
| **Size** | -0.164 | 0.005 | -36.167 | 2.4e-129 |
| **Size \* Is Ensemble** | -0.281 | 0.014 | -19.696 | 3.5e-61 |

Full ensemble specific results and OLS model details can be found in the supplementary material at sahahn.github.io/parc_scaling/ensemble_comparison, and full grid vs. random at sahahn.github.io/parc_scaling/grid_vs_random.

### Highest Performance

The subsets of results plotted in Figure 6 were chosen to highlight the best performing individual results as identified throughout the previous analyses, as well as introduce the results from the special ‘All’ ensemble. In summary, the SVM outperformed other pipelines in the inter-pipeline comparison (Figure 3) and existing parcellations outperformed randomly generated ones (Figure 2), which is why the subset ‘SVM Non-Random Existing’ was included. Similarly, an inter-pipeline comparison was conducted for the ensemble based results, which suggested the SVM based ensembles had better performance relative to the other pipelines beyond size 500 (see sahahn.github.io/parc_scaling/ensemble_by_pipeline). Last, we included the ‘All’ ensemble, which as expected performed as well or better than the SVM based ensemble (as it has all of the same base ensembled pipelines). Combined, these subsets of results highlight the best performing strategies across different spatial scales.

**Figure 6.**
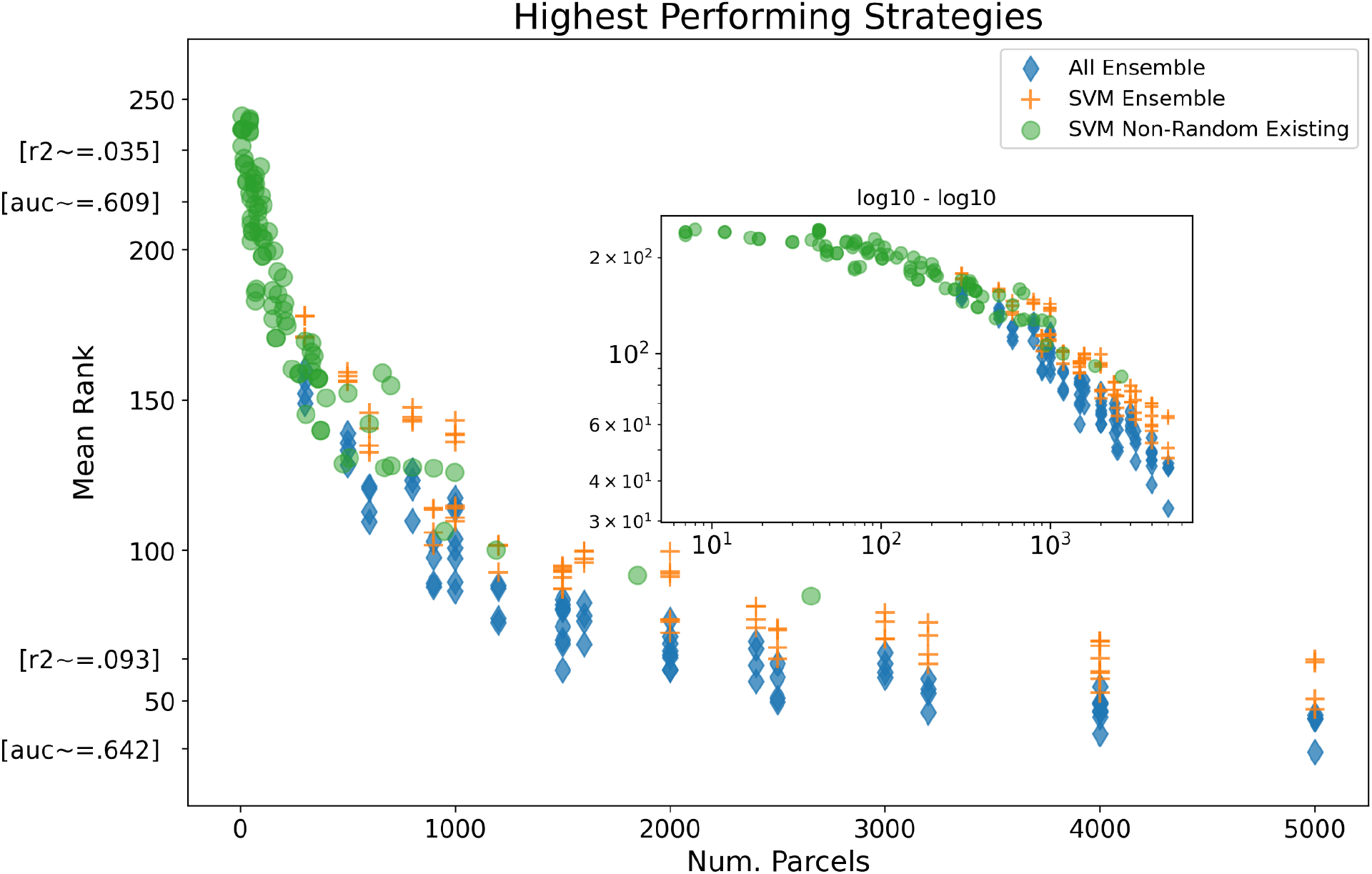
plots the mean ranks as computed between a specially chosen set of high performing parcellation-pipeline results. This subset includes foremost results from the ‘All’ ensemble (blue diamond), where a single ensemble is used to combine predictions from Elastic-Net, LGBM and SVM pipelines all trained on the same set of fixed-size parcellations (across size parcellation ensemble results not included here). Further, results from just the fixed size SVM ensemble are shown (orange plus), as well as the single, non randomly generated, parcellation results from just the SVM based pipeline (green circle). Results are plotted by the number of parcels on the x-axis, along with a log10-log10 inset version of the same plot. Example estimates of mean R2 and ROC AUC are also provided on the y-axis as estimated across only the regression or binary target variables from parcellation results as close as possible to that rank.

## Discussion

The current study tested the impact of parcellation choice as a factor in out of sample performance across a variety of target variables using structural MRI data. To do this we estimated the 5-fold predictive performance across different combinations of parcellation and ML pipeline for all target variables. Across all combinations, we found that, in general, parcellations with more parcels performed better than those with fewer parcels up to about 4,000 parcels, although at the expense of greater variability in performance. The increase in performance as the number of parcels increases can be well characterized by a power law scaling relationship (i.e., diminishing returns to gains in performance as resolution increases). We also found that pre-existing parcellations based on cytoarchitecture or functional MRI performed better than randomly derived parcellations of comparable size, suggesting merit to existing parcellation schemes. However, pre-existing parcellations tended to have fewer parcels than our results suggest will perform best. Among the ML pipelines and strategies tested, SVM based pipelines had the best predictive performance, although this could be further improved by ensembling over multiple parcellations and/or pipelines.

One of the key goals of this work was describing the relationship between spatial scale (i.e., number of parcels) and predictive performance. Based on our results, this relationship appears to follow a power-law scaling, although the details of this scaling can vary with choice of pipeline and parcellation strategy. We identified scaling across roughly 3 orders of magnitude (∼10-4000) with coefficients between ¼ and ¾ (exact coefficient dependent on specific setup). We further tested how stable this relationship was when compared in an intra-pipeline fashion, finding that the general pattern was preserved. That said, the SVM-based results varied significantly from the other two pipelines, with a steeper estimated scaling coefficient and a larger estimated region of scaling.

There are several possible reasons for the increase in predictive performance with parcellation resolution. First, we note that by ‘applying’ parcellations within the context of the analyses above, we are simply taking the mean value across a contiguous region of vertices. That means that when the region of interest is centered on a non-homogeneous area that the mean value of the region will essentially have added noise. Therefore by increasing the resolution of the parcellation, with a larger number of smaller parcels, each parcel will be less likely to span across multiple distinct structures (i.e., “true regions”). That said, once the resolution becomes too fine grained, subdividing true regions of interest can also introduce noisy estimates.

The very existence of a tightly knit scaling between parcellation granularity and out-of-sample phenotypic predictive performance may itself be meaningful, regardless of the exact scaling or range of scaling. Because existing parcellations have a relatively small number of parcels, this scaling suggests that commonly employed parcellations may be too coarse to capture some important inter-individual phenotypic variance. In other words, up to a certain resolution there is still valuable information lost due to averaging over too large an area, potentially containing distinct functions. This increasing marginal utility of smaller and smaller brain regions is fully compatible with the view of the brain as a “hierarchically organized system”, where meaningful cortical areas can be composed at multiple resolutions (Eickhoff 2018). In this sense, as the number of parcels increases it may provide a better mapping onto different resolutions of “true” cortical areas. For example, a small number of parcels may only be capturing differences at the highest hierarchical level, but as more parcels are added the gains in performance we found may represent a mapping onto the next meaningful hierarchical resolution of cortical areas.

The observed scaling between performance and size can also be used to inform practical recommendations for researchers. First, we note that current common practices with respect to choice of parcellation may not fully be exploiting all available predictive information. As a representative case, we focus on the results from the two FreeSurfer extracted sets of ROIs. As shown in Figure 2, we note that the observed mean rank for both FreeSurfer based parcellations are almost fully explained by their number of parcels (Desikan - 70, Destrieux - 150) and their status as an existing parcellation. For these and other existing parcellations commonly used, their inability to exploit all predictive information therefore comes down primarily to the number of unique ROI’s. Therefore, when prioritizing predictive capability, a simple heuristic is to select the highest possible resolution parcellation, in this case Destrieux over Desikan. Importantly, the diminishing returns nature of the scaling relationship (e.g., consider relative performance gains between sizes 100 to 3000 vs. 3000 to 4000) as well as the region where scaling ends (e.g., 20,000 parcels likely performs worse than 3000), should also be taken into account.

Notably, it is not necessarily true that parcellations with a higher number of parcels will always perform better. For example, comparing between randomly generated parcellations and existing literature-based parcellations revealed consistently better performance for existing parcellations. This could suggest that, on average, the existing parcellations map better onto meaningful neuroanatomy relative to random parcellations of similar size. Different phenotypes of interest also vary in how much they follow the observed scaling relationship. We also found an interesting increase in spread of mean ranks as the number of parcels grew, where not only the mean rank increased but also the inter quartile range at each size increased. This behavior is likely a result of the distributed and complex brain-based nature of the phenotypes studied, where different targets may have different optimal resolutions. That said, the pattern in the average case remains clear and, we argue, is still meaningful despite recognizing that variation exists across possible phenotypes.

We also explored the influence of choice of ML pipeline. For parcellations larger than size ∼100 the SVM based pipeline outperformed all other pipelines, with the Elastic-Net the best at smaller sizes. The LGBM tree-based pipeline was not competitive at any size, an observation in line with recent work based on sMRI data from UK Biobank participants (Schulz 2020). We further ruled out the hypothesis that the SVM’s comparative scaling was driven by unique access to a nested front end feature selection (See sahahn.github.io/parc_scaling/effect_of_fs.html). While perhaps interesting conceptually, treating choice of parcellation as a nested hyper-parameter, in practice, yielded lackluster results, especially when compared with the ensemble based methods. We observed that this approach fell closely in line with expected random parcellation performance at the same size. In contrast, we observed a significant performance improvement from the multiple parcellation ensemble-based strategies when compared to the single parcellation only results. Notably, the ensemble-based random parcellations continue to exhibit scaling beyond the ∼4000 range where scaling was estimated to have ended with respect to the analyses with single parcellations. These results establish the merit in constructing ensembles across multiple parcellations to achieve maximal predictive performance. Specifically, we found no significant differences in predictive performance between the voting and stacking ensemble approaches tested. We did observe significant differences between ensembles with random parcellations of the same size versus ensembles with parcellations of multiple sizes, in this case finding that the fixed size parcellations on average performed better. Therefore, to maximize predictive performance and computational demands, we recommend that, of the ensemble methods tested, fixed size parcellations with a voting ensemble be used in future work.

One obvious potential explanation for the observed ensemble performance gain is that it is due solely to an inherent utility of ensembling, which has been shown to reliably increase performance across a wide range of ML applications (Dietterich 2000, Zhou 2009). On the other hand, ensembles as employed in this work are specifically designed to capture information from multiple overlapping parcellations. It is plausible that the performance boost obtained by this methodology may be related to the boost from increasing resolution; this could indicate that the “true” best parcellations are not uniform and universal. Instead, by allowing overlapping parcellations, more predictive information can be extracted despite noisy ground truth data. Alternatively, it could also be that ensembling over multiple views or sizes provides benefit by forcing different classifiers to exploit different unique predictive signals (Allen-Zhu 2020). The cortical surface exhibits high covariance between different brain regions on measures employed as input features (e.g., cortical thickness); it may therefore be reasonable to assume that there is more than one multivariate predictive pattern capable of performing well out of sample on the target of interest (Alexander-Bloch, 2013). If this is the case, then different instances of random parcellations may help base estimators of the ensemble learn distinct predictive patterns that when combined can improve predictive power.

Throughout this work, different strategies and algorithms outperformed others at different spatial scales. The two top performing strategies were the SVM based pipeline with existing parcellations and an ensemble combining the three ML algorithms (the ‘All ensemble) with randomly generated parcellations of the same size. The ‘All’ ensemble results in particular were interesting as they were both the most complex, where predictions were averaged across multiple combinations of pipeline and parcellation, but were also consistently the highest performing. Despite the noted variability across targets and noting that one parcellation / algorithm will never *always* provide the best performance, we still believe it useful to characterize the general patterns which influence performance (Wolpert 1995). In this case, we draw attention to the relative merit of SVM based pipelines, non-random individual parcellations, and ensembling across multiple parcellations and / or ML algorithms.

In practice, a researcher’s choice of parcellation often does not solely prioritize predictive performance. Instead other key influences include interpretability and computational resources. As parcellation resolution increases, the number of regions to interpret as well as runtime for the analysis will scale accordingly. Runtime and memory requirements are also influenced greatly by choice of algorithm. For example, on a machine with 8 cores and 8GB of RAM, an iteration of the 5-fold cross-validation with Elastic-Net and 200 parcels required around 15 minutes to run compared to an SVM and 2000 parcels which required around 15 hours to run. Likewise, evaluations can quickly grow intractable; for example, on a machine with 4 cores and 256GB of RAM, employing non parcellated vertex-level data directly (59,412 “parcels”) required over a week to finish even a single of the five folds. Another inherent benefit to employing some existing parcellations, regardless of their number of parcels, is their popularity itself, which can facilitate easier comparisons via named regions with prior published work. Trade-offs are discussed in further detail at sahahn.github.io/parc_scaling/trade_offs.html.

Extensions of the present study beyond structural MRI to other MRI modalities represents a potentially interesting future direction, especially in the case of functional connectome data where features grow exponentially with number of regions. Likewise, it is unknown if the current findings apply to segmentation based on volumetric data. Other possible extensions could be to investigate further how results change if each structural metric (i.e., thickness, surface area, etc) were treated independently, with for example ensembles over structural metric specific ML pipelines. Another factor is that neuroimaging based ML has been shown to exhibit a performance to sample size scaling, where larger samples, up to a certain point, can continue to yield performance boosts (Schulz 2020). As such, it is important to note that the influence of sample size on the observed parcellation performance scaling remains unknown. As the current work uses samples of around 7,500 in each fold, it is not clear from current results how well the observed performance-size scaling will hold for smaller studies. Another future direction could be to test different ML estimators, for example deep learning based algorithms that may exhibit different scaling, as they can in theory better handle data with structured high dimensional feature spaces (He 2020, Abrol 2021).

## Conclusion

In testing a variety of parcellation schemes and ML modeling approaches, we have identified an apparent power law scaling of increasing predictive performance by increasing parcellation resolution. The details of this relationship were found to vary according to type of parcellation as well as ML pipeline employed, although the general pattern proved stable. The large sample size, range of predictive targets, and collection of existing and random parcellations tested all serve to lend confidence to the observed results. Researchers selecting a parcellation for predictive modelling may wish to consider this size-performance trade-off in addition to other factors such as interpretability and computational resources. We also highlighted important factors that improved performance above and beyond the size-scaling, for example, finding that existing parcellations performed better than randomly generated parcellations. Further, we demonstrated the benefit of ensembling over multiple parcellations, which yielded a performance boost relative to results from single parcellations.

## Acknowledgments

Sage Hahn, Max M. Owens, DeKang Yuan and Anthony C Juliano were supported by NIDA grant T32DA043593.

Data used in the preparation of this article were obtained from the ABCD Study (https://abcdstudy.org) held in the NDA. This is a multisite, longitudinal study designed to recruit more than 10,000 children ages 9–10 years old and follow them over 10 years into early adulthood. The ABCD study is supported by the National Institutes of Health and additional federal partners under award numbers U01DA041048, U01DA050989, U01DA051016, U01DA041022, U01DA051018, U01DA051037, U01DA050987, U01DA041174, U01DA041106, U01DA041117, U01DA041028, U01DA041134, U01DA050988, U01DA051039, U01DA041156, U01DA041025, U01DA041120, U01DA051038, U01DA041148, U01DA041093, U01DA041089, U24DA041123 and U24DA041147. A full list of supporters is available at https://abcdstudy.org/federal-partners.html. A listing of participating sites and a complete listing of the study investigators can be found at https://abcdstudy.org/consortium_members/.

Computations were performed on the Vermont Advanced Computing Core supported, in part, by NSF award number OAC-1827314.

We would also like to thank the other members of the Hugh Garavan lab for their support throughout this project.

## References

Abrol, A., Fu, Z., Salman, M., Silva, R., Du, Y., Plis, S., & Calhoun, V. (2021). Deep learning encodes robust discriminative neuroimaging representations to outperform standard machine learning. Nature communications, 12(1), 1–17.

Alexander-Bloch, A., Raznahan, A., Bullmore, E., & Giedd, J. (2013). The convergence of maturational change and structural covariance in human cortical networks. Journal of Neuroscience, 33(7), 2889–2899.

Allen, E. A., Erhardt, E. B., Damaraju, E., Gruner, W., Segall, J. M., Silva, R. F., … & Calhoun, V. D. (2011). A baseline for the multivariate comparison of resting-state networks. Frontiers in systems neuroscience, 5, 2.

Allen-Zhu, Z., & Li, Y. (2020). Towards Understanding Ensemble, Knowledge Distillation and Self-Distillation in Deep Learning. arXiv preprint 2012.09816.

Amari, S. I., & Wu, S. (1999). Improving support vector machine classifiers by modifying kernel functions. Neural Networks, 12(6), 783–789.

Arslan, S., Ktena, S. I., Makropoulos, A., Robinson, E. C., Rueckert, D., & Parisot, S. (2018). Human brain mapping: A systematic comparison of parcellation methods for the human cerebral cortex. Neuroimage, 170, 5–30.

Baldassano, C., Beck, D. M., & Fei-Fei, L. (2015). Parcellating connectivity in spatial maps. PeerJ, 3, e784.

Bellec, P. (2013, June). Mining the hierarchy of resting-state brain networks: selection of representative clusters in a multiscale structure. In 2013 International Workshop on Pattern Recognition in Neuroimaging (pp. 54–57). IEEE.

Bhagwat, N., Pipitone, J., Voineskos, A. N., Chakravarty, M. M., & Alzheimer’s Disease Neuroimaging Initiative. (2019). An artificial neural network model for clinical score prediction in Alzheimer disease using structural neuroimaging measures. Journal of psychiatry & neuroscience: JPN, 44(4), 246.

Boeke, E. A., Holmes, A. J., & Phelps, E. A. (2020). Toward robust anxiety biomarkers: a machine learning approach in a large-scale sample. Biological Psychiatry: Cognitive Neuroscience and Neuroimaging, 5(8), 799–807.

Brodmann, K. (1909). Vergleichende Lokalisationslehre der Grosshirnrinde in ihren Prinzipien dargestellt auf Grund des Zellenbaues. Barth.

Challenges. ”NeuroImage, vol. 197, 2019, pp. 652–656., doi:10.1016/j.neuroimage.2018.10.003.

Chen, T., He, T., Benesty, M., Khotilovich, V., & Tang, Y. (2015). Xgboost: extreme gradient boosting. R package version 0.4-2, 1–4.

Clauset, A., Young, M., & Gleditsch, K. S. (2007). On the frequency of severe terrorist events. Journal of Conflict Resolution, 51(1), 58–87.

Craddock, C., Sikka, S., Cheung, B., Khanuja, R., Ghosh, S. S., Yan, C., … & Milham, M. (2013). Towards automated analysis of connectomes: The configurable pipeline for the analysis of connectomes (c-pac). Front Neuroinform, 42.

Craddock, R. C., James, G. A., Holtzheimer III, P. E., Hu, X. P., & Mayberg, H. S. (2012). A whole brain fMRI atlas generated via spatially constrained spectral clustering. Human brain mapping, 33(8), 1914–1928.

Dadi, K., Rahim, M., Abraham, A., Chyzhyk, D., Milham, M., Thirion, B., … & Alzheimer’s Disease Neuroimaging Initiative. (2019). Benchmarking functional connectome-based predictive models for resting-state fMRI. Neuroimage, 192, 115–134.

Dadi, K., Varoquaux, G., Machlouzarides-Shalit, A., Gorgolewski, K. J., Wassermann, D., Thirion, B., and Mensch, A. (2020). Fine-grain atlases of functional modes for fMRI analysis. NeuroImage.

Davatzikos, C. (2019). Machine learning in neuroimaging: Progress and challenges. NeuroImage, 197, 652.

Desikan, R. S., Ségonne, F., Fischl, B., Quinn, B. T., Dickerson, B. C., Blacker, D., … & Killiany, R. J. (2006). An automated labeling system for subdividing the human cerebral cortex on MRI scans into gyral based regions of interest. Neuroimage, 31(3), 968–980.

Destrieux, C., Fischl, B., Dale, A., & Halgren, E. (2010). Automatic parcellation of human cortical gyri and sulci using standard anatomical nomenclature. Neuroimage, 53(1), 1–15.

Dietterich, T. G. (2000, June). Ensemble methods in machine learning. In International workshop on multiple classifier systems (pp. 1–15). Springer, Berlin, Heidelberg.

Eickhoff, S. B., Constable, R. T., & Yeo, B. T. (2018). Topographic organization of the cerebral cortex and brain cartography. Neuroimage, 170, 332–347.

Eickhoff, S. B., Stephan, K. E., Mohlberg, H., Grefkes, C., Fink, G. R., Amunts, K., & Zilles, K. (2005). A new SPM toolbox for combining probabilistic cytoarchitectonic maps and functional imaging data. Neuroimage, 25(4), 1325–1335.

Fan, L., Li, H., Zhuo, J., Zhang, Y., Wang, J., Chen, L., … & Jiang, T. (2016). The human brainnetome atlas: a new brain atlas based on connectional architecture. Cerebral cortex, 26(8), 3508–3526.

Fischl, B. (2012). FreeSurfer. Neuroimage, 62(2), 774–781.

Franke, K., Ziegler, G., Klöppel, S., Gaser, C., & Alzheimer’s Disease Neuroimaging Initiative. (2010). Estimating the age of healthy subjects from T1-weighted MRI scans using kernel methods: exploring the influence of various parameters. Neuroimage, 50(3), 883–892.

Glasser, M. F., Coalson, T. S., Robinson, E. C., Hacker, C. D., Harwell, J., Yacoub, E., … & Van Essen, D. C. (2016). A multi-modal parcellation of human cerebral cortex. Nature, 536(7615), 171–178.

Glasser, M. F., Sotiropoulos, S. N., Wilson, J. A., Coalson, T. S., Fischl, B., Andersson, J. L., … & Van Essen, D. C. (2013). The minimal preprocessing pipelines for the Human Connectome Project. Neuroimage, 80, 105–124.

Gordon, E. M., Laumann, T. O., Adeyemo, B., Huckins, J. F., Kelley, W. M., & Petersen, S. E. (2016). Generation and evaluation of a cortical area parcellation from resting-state correlations. Cerebral cortex, 26(1), 288–303.

Gowin, J. L., Ernst, M., Ball, T., May, A. C., Sloan, M. E., Tapert, S. F., & Paulus, M. P. (2019). Using neuroimaging to predict relapse in stimulant dependence: A comparison of linear and machine learning models. NeuroImage: Clinical, 21, 101676.

Hagler Jr, D. J., Hatton, S., Cornejo, M. D., Makowski, C., Fair, D. A., Dick, A. S., … & Dale, A. M. (2019). Image processing and analysis methods for the Adolescent Brain Cognitive Development Study. Neuroimage, 202, 116091.

Hahn, S, Mackey, S, Cousijn, J, et al. (2020) Predicting alcohol dependence from multi-site brain structural measures. Hum Brain Mapp. 2020; 1–11. https://doi.org/10.1002/hbm.25248

Hahn, S., Yuan, D. K., Thompson, W. K., Owens, M., Allgaier, N., & Garavan, H. (2021). Brain Predictability toolbox: A Python library for neuroimaging-based machine learning. Bioinformatics (Oxford, England).

Hammers, A., Allom, R., Koepp, M. J., Free, S. L., Myers, R., Lemieux, L., … & Duncan, J. S. (2003). Three-dimensional maximum probability atlas of the human brain, with particular reference to the temporal lobe. Human brain mapping, 19(4), 224–247.

Harrison, S. J., Woolrich, M. W., Robinson, E. C., Glasser, M. F., Beckmann, C. F., Jenkinson, M., & Smith, S. M. (2015). Large-scale probabilistic functional modes from resting state fMRI. NeuroImage, 109, 217–231.

He, N., Rolls, E. T., Zhao, W., & Guo, S. (2020). Predicting human inhibitory control from brain structural MRI. Brain imaging and behavior, 14(6), 2148–2158.

He, T., Kong, R., Holmes, A. J., Nguyen, M., Sabuncu, M. R., Eickhoff, S. B., … & Yeo, B. T. (2020). Deep neural networks and kernel regression achieve comparable accuracies for functional connectivity prediction of behavior and demographics. NeuroImage, 206, 116276.

Hong, S., Liu, Y. S., Cao, B., Cao, J., Ai, M., Chen, J., … & Kuang, L. (2020). Identification of suicidality in adolescent major depressive disorder patients using sMRI: A machine learning approach. Journal of Affective Disorders.

Janssen, R. J., Mourão-Miranda, J., & Schnack, H. G. (2018). Making individual prognoses in psychiatry using neuroimaging and machine learning. Biological Psychiatry: Cognitive Neuroscience and Neuroimaging, 3(9), 798–808.

Jenkinson, M., Beckmann, C. F., Behrens, T. E., Woolrich, M. W., & Smith, S. M. (2012). Fsl. Neuroimage, 62(2), 782–790.

Joliot, M., Jobard, G., Naveau, M., Delcroix, N., Petit, L., Zago, L., … & Tzourio-Mazoyer, N. (2015). AICHA: An atlas of intrinsic connectivity of homotopic areas. Journal of neuroscience methods, 254, 46–59.

Jollans, L., Boyle, R., Artiges, E., Banaschewski, T., Desrivières, S., Grigis, A., … & Schumann, G. (2019). Quantifying performance of machine learning methods for neuroimaging data. NeuroImage, 199, 351–365.

Karch, J. D., Filevich, E., Wenger, E., Lisofsky, N., Becker, M., Butler, O., … & Kühn, S. (2019). Identifying predictors of within-person variance in MRI-based brain volume estimates. NeuroImage, 200, 575–589.

Khundrakpam, B. S., Tohka, J., Evans, A. C., & Brain Development Cooperative Group. (2015). Prediction of brain maturity based on cortical thickness at different spatial resolutions. Neuroimage, 111, 350–359.

Marcus, D., Harwell, J., Olsen, T., Hodge, M., Glasser, M., Prior, F., … & Van Essen, D. (2011). Informatics and data mining tools and strategies for the human connectome project. Frontiers in neuroinformatics, 5, 4.

Mateos-Pérez, J. M., Dadar, M., Lacalle-Aurioles, M., Iturria-Medina, Y., Zeighami, Y., & Evans, A. C. (2018). Structural neuroimaging as clinical predictor: A review of machine learning applications. NeuroImage: Clinical, 20, 506–522.

Mwangi, B., Tian, T. S., & Soares, J. C. (2014). A review of feature reduction techniques in neuroimaging. Neuroinformatics, 12(2), 229–244.

Nielsen, A. N., Barch, D. M., Petersen, S. E., Schlaggar, B. L., & Greene, D. J. (2019). Machine learning with neuroimaging: evaluating its applications in psychiatry. Biological Psychiatry: Cognitive Neuroscience and Neuroimaging.

Owens, M. M., Potter, A., Hyatt, C., Albaugh, M., Thompson, W. K., Jernigan, T., … & Garavan, H. (2020). Recalibrating expectations about effect size: A multi-method survey of effect sizes in the ABCD study.

Pedregosa, F., Varoquaux, G., Gramfort, A., Michel, V., Thirion, B., Grisel, O., … & Vanderplas, J. (2011). Scikit-learn: Machine learning in Python. the Journal of machine Learning research, 12, 2825–2830.

Power, J. D., Cohen, A. L., Nelson, S. M., Wig, G. S., Barnes, K. A., Church, J. A., … & Petersen, S. E. (2011). Functional network organization of the human brain. Neuron, 72(4), 665–678.

Rapin, J., & Teytaud, O. (2018). Nevergrad-A gradient-free optimization platform. version 0.2. 0, https://GitHub.com/FacebookResearch/Nevergrad.

Sabuncu, M. R., Desikan, R. S., Sepulcre, J., Yeo, B. T. T., Liu, H., Schmansky, N. J., … & Alzheimer’s Disease Neuroimaging Initiative. (2011). The dynamics of cortical and hippocampal atrophy in Alzheimer disease. Archives of neurology, 68(8), 1040–1048.

Salehi, M., Greene, A. S., Karbasi, A., Shen, X., Scheinost, D., & Constable, R. T. (2020). There is no single functional atlas even for a single individual: Functional parcel definitions change with task. NeuroImage, 208, 116366.

Sato, J. R., Hoexter, M. Q., de Magalhães Oliveira Jr, P. P., Brammer, M. J., Murphy, D., Ecker, C., & MRC AIMS Consortium. (2013). Inter-regional cortical thickness correlations are associated with autistic symptoms: a machine-learning approach. Journal of psychiatric research, 47(4), 453–459.

Schaefer, A., Kong, R., Gordon, E. M., Laumann, T. O., Zuo, X. N., Holmes, A. J., … & Yeo, B. T. (2018). Local-global parcellation of the human cerebral cortex from intrinsic functional connectivity MRI. Cerebral cortex, 28(9), 3095–3114.

Schulz, M. A., Yeo, B. T., Vogelstein, J. T., Mourao-Miranada, J., Kather, J. N., Kording, K., … & Bzdok, D. (2020). Different scaling of linear models and deep learning in UKBiobank brain images versus machine-learning datasets. Nature communications, 11(1), 1–15.

Seabold, Skipper, and Josef Perktold. “statsmodels: Econometric and statistical modeling with pythonx.“ Proceedings of the 9th Python in Science Conference. 2010.

Shen, X., Tokoglu, F., Papademetris, X., & Constable, R. T. (2013). Groupwise whole-brain parcellation from resting-state fMRI data for network node identification. Neuroimage, 82, 403–415.

Squarcina, L., Castellani, U., Bellani, M., Perlini, C., Lasalvia, A., Dusi, N., … & Alessandrini, F. (2017). Classification of first-episode psychosis in a large cohort of patients using support vector machine and multiple kernel learning techniques. NeuroImage, 145, 238–245.

Sripada, C. S., Kessler, D., & Angstadt, M. (2014). Lag in maturation of the brain’s intrinsic functional architecture in attention-deficit/hyperactivity disorder. Proceedings of the National Academy of Sciences, 111(39), 14259–14264.

Tzourio-Mazoyer, N., Landeau, B., Papathanassiou, D., Crivello, F., Etard, O., Delcroix, N., … & Joliot, M. (2002). Automated anatomical labeling of activations in SPM using a macroscopic anatomical parcellation of the MNI MRI single-subject brain. Neuroimage, 15(1), 273–289.

Urchs, S., Armoza, J., Moreau, C., Benhajali, Y., St-Aubin, J., Orban, P., & Bellec, P. (2019). MIST: A multi-resolution parcellation of functional brain networks. MNI Open Research, 1(3), 3.

Van Essen, D. C., Glasser, M. F., Dierker, D. L., Harwell, J., & Coalson, T. (2012). Parcellations and hemispheric asymmetries of human cerebral cortex analyzed on surface-based atlases. Cerebral cortex, 22(10), 2241–2262.

Varoquaux, G., Gramfort, A., Pedregosa, F., Michel, V., & Thirion, B. (2011, July). Multi-subject dictionary learning to segment an atlas of brain spontaneous activity. In Biennial International Conference on information processing in medical imaging (pp. 562–573). Springer, Berlin, Heidelberg.

Von Luxburg, U., & Schölkopf, B. (2011). Statistical learning theory: Models, concepts, and results. In Handbook of the History of Logic (Vol. 10, pp. 651–706).

North-Holland Wang, L., Mruczek, R. E., Arcaro, M. J., & Kastner, S. (2015). Probabilistic maps of visual topography in human cortex. Cerebral cortex, 25(10), 3911–3931.

Whitaker, K. J., Vértes, P. E., Romero-Garcia, R., Váša, F., Moutoussis, M., Prabhu, G., … & NSPN Consortium. (2016). Adolescence is associated with genomically patterned consolidation of the hubs of the human brain connectome. Proceedings of the National Academy of Sciences, 113(32), 9105–9110.

Wolpert, D. H. (1992). Stacked generalization. Neural networks, 5(2), 241–259.

Wolpert, D. H., & Macready, W. G. (1995). No free lunch theorems for search (Vol. 10). Technical Report SFI-TR-95-02-010, Santa Fe Institute.

Wu, J., Ngo, G. H., Greve, D., Li, J., He, T., Fischl, B., … & Yeo, B. T. (2018). Accurate nonlinear mapping between MNI volumetric and FreeSurfer surface coordinate systems. Human brain mapping, 39(9), 3793–3808.

Yeo, B. T., Krienen, F. M., Chee, M. W., & Buckner, R. L. (2014). Estimates of segregation and overlap of functional connectivity networks in the human cerebral cortex. Neuroimage, 88, 212–227.

Zhou, Z. H. (2009). Ensemble learning. Encyclopedia of biometrics, 1, 270–273.

de Wit, S., Ziermans, T. B., Nieuwenhuis, M., Schothorst, P. F., van Engeland, H., Kahn, R. S., … & Schnack, H. G. (2017). Individual prediction of long-term outcome in adolescents at ultra-high risk for psychosis: Applying machine learning techniques to brain imaging data. Human brain mapping, 38(2), 704–714.

von Economo, C. F., & Koskinas, G. N. (1925). Die cytoarchitektonik der hirnrinde des erwachsenen menschen. J. Springer.

